# Limits and promises of Earth Observation Foundation Models in Predicting Multi-Trophic Soil Biodiversity

**DOI:** 10.1101/2025.05.16.654504

**Authors:** Selene Cerna, Sara Si-Moussi, Irene Calderón-Sanou, Vincent Miele, Wilfried Thuiller

## Abstract

Soil biodiversity is essential for terrestrial ecosystems, influencing nutrient cycling, carbon sequestration, agricultural productivity, and resilience to environmental changes. Yet, it faces significant threats from land-use changes, pollution, agricultural intensification, and climate change. Effective predictive modeling tools are urgently needed to inform conservation and management strategies. Although species distribution models (SDMs) have been successful for aboveground biodiversity, their application to soil biodiversity is limited by scarce large-scale datasets with spatial mismatches between environmental data and soil habitats. Recent advances, including environmental DNA (eDNA) metabarcoding, now allow extensive multi-taxa assessments of soil biodiversity. Simultaneously, remote sensing technologies provide high-resolution spatial data, potentially overcoming traditional coarse-gridded environmental limitations. This study evaluates Earth Observation Foundation (EOF) models, deep learning models pretrained on massive remote sensing datasets to summarize earth observation images into embeddings, to predict multi-trophic soil biodiversity in the French Alps. We compare models using EOF-derived embeddings from orthophotos with coarse-gridded and high-quality in-situ variables. We modeled relative abundance for 51 trophic groups across seven taxa using Random Forest, Light Gradient Boosting Machine, and Artificial Neural Networks, evaluating four data configurations: coarse-gridded environmental data, high-quality in-situ data, EOF embeddings, and a hybrid embedding-tabular approach. High-quality in-situ climate and soil data consistently delivered the highest predictive accuracy, especially for microbial and fungal groups. EOF embeddings provided valuable spatial context but did not surpass in-situ data performance, showing partial redundancy. Integrating remote sensing data can enhance biodiversity modeling in areas lacking detailed in-situ measurements, underscoring their complementary role in ecological assessments.

## 1. Introduction

Soil biodiversity is central to the functioning of terrestrial ecosystems [1]. It is the cornerstone of processes that sustain life on Earth, such as nutrient cycling, carbon sequestration, and resilience to environmental changes [2]. It represents approximately 59% of global biodiversity [3], encompassing organisms from microorganisms and microfauna, responsible for converting inorganic and organic compounds into plant-accessible forms, to mesofauna and macrofauna, which facilitate nutrient cycling and enhance soil physical structure and water retention [4, 5]. These diverse biological processes collectively maintain soil fertility, support agricultural productivity, and underpin ecosystem services worldwide.

Despite its crucial role, soil biodiversity is under increasing threat due to human activities and environmental change [6]. Land use transformations and agricultural intensification disrupt soil habitats, desertification exhausts critical nutrients, pollution degrades soil quality, and climate change imposes additional temperature and moisture constraints [7, 8, 9, 10]. These threats disrupt soil habitats, degrade soil quality, and impose additional constraints on biodiversity, highlighting the urgent need for effective tools capable of accurately predicting soil biodiversity patterns to inform conservation strategies, sustainable agricultural practices, and ecosystem management [11].

Species distribution models (SDMs) are currently the primary tools employed to predict biodiversity patterns [12]. These are computational tools that relate observed species (or groups of species) occurrence, abundance or biomass data to selected environmental variables [13]. These variables usually take the form of tabular climatic factors, soil conditions, vegetation cover, land use, or even other organisms [14]. These variables are called tabular because they provide a single measure for a single sampling unit (e.g., mean temperature, soil pH). Although successful in modeling above-ground taxa such as plants [15, 16], vertebrates [17] or invertebrates [18], their application to soil biodiversity has remained limited (but see [19, 20, 21]), primarily due to two main constraints. First, high-quality, large-scale datasets of soil biodiversity have historically been scarce, severely restricting model development and validation. The recent advent of environmental DNA (eDNA) metabarcoding now addresses this critical data limitation by enabling large-scale, multi-taxa soil biodiversity assessments, including taxa that were poorly documented before such as bacteria, fungi, and microfauna [22]. Second, there is a significant spatial mismatch between available environmental data and the scales at which soil organisms experience their environment. In-situ or highly resolved climate or soil variables are usually good predictors of soil biodiversity [23], but they usually do not allow predictions in non-sampled areas which are especially important for mapping and conservation purposes. Instead, most global or regional environmental datasets are available in relatively coarse resolutions (1 km²), predominantly representing aboveground conditions and insufficiently capturing critical soil-specific drivers such as soil pH, organic matter, or fine-scale vegetation heterogeneity ([24, 25]). Remote sensing technologies offer a promising solution, as satellite imagery (e.g., Sentinel-2 at 10m resolution), aerial RGB-IR photographs, and LiDAR data provide spatially explicit, high-resolution information on vegetation structure, landscape heterogeneity indirectly relevant to below-ground conditions [26, 27]. Recent studies have shown the potential of hyperspectral data to predict microbial diversity patterns at broad scales [28]. However, the explicit use and comparative evaluation of remote sensing data versus high-quality in-situ measurements or coarse-gridded environmental variables for predicting multi-trophic soil biodiversity remains largely unexplored.

Moreover, remote sensing data, particularly aerial images, inherently offer richer spatial context beyond single-point measurements, capturing landscape structure and spatial context that can be integrated in predictive models. Deep learning approaches, specifically Deep-Species Distribution Models (DeepSDMs, [29]), effectively leverage this embedded spatial structure by transforming the remote sensing data into efficient predictors. Neural networks excel at representation learning, enabling them to transform complex remote sensing image data into compact vector representations known as ”embeddings”. This approach bypasses the challenges posed by the high dimensionality and complexity of such data. However, training deep models for this task presents significant challenges, particularly due to the need for massive datasets and substantial computing power. These challenges are even more pronounced in soil biodiversity studies, where datasets tend to be relatively small.

Earth Observation Foundation (EOF) models, leveraging self-supervised pre-training on massive unlabeled remote sensing datasets, offer great potential in overcoming these limitations by generating general-purpose representations easily fine-tuned for specialized ecological applications. These models can handle satellite imagery, environmental data, and socio-economic factors to better understand phenomena like land-cover changes and biodiversity loss [30]. Notable examples include *DiffusionSat* [31], which excels in various generative tasks from super-resolution to temporal image generation; *DOFA*[32], a dynamic one-for-all model inspired by neural plasticity to handle diverse sensors; *SatDINO* [33] (for convenience, we refer to the EOF model described in [33] as ’SatDINO’ in this study), which leverages high-resolution aerial RGB imagery, originally trained to predict canopy height from LiDAR, with Vision Transformer-based encoders in a unsupervised manner; and *Prithvi-EO-2.0* [34], a multi-temporal foundation model trained on Landsat and Sentinel-2 data for land-use classification and ecosystem monitoring. Despite these breakthroughs, EOF models have not yet been rigorously tested for biodiversity prediction, nor systematically compared against conventional gridded-environmental datasets or detailed in-situ measurements. Since they alleviate the issue of data limitation for small datasets, they thus offer the possibility of training soil biodiversity SDMs while taking advantage of the breadth of information contained in images. Whether this provide much as quality data as that in-situ measurements remain to be tested.

To address this critical research gap, we systematically evaluate the efficiency of EOF models in predicting soil biodiversity, specifically focusing on multi-trophic groups identified through eDNA metabarcoding data. Using data from the ORCHAMP observatory [13], we predict the relative abundance of 51 trophic groups across seven major biological categories: Bacteria, Fungi, Protist, Oligochaete, Insect, Collembola, and Metazoa. For clarity, we refer to Metazoa here as the remaining animal taxa excluding Oligochaete, Insect, and Collembola, which are treated as separate groups due to their distinct ecological roles and their associated specific DNA markers. Together, these trophic groups span a wide range of ecological functions, collectively illustrating the complexity of soil life [35]. Specifically, we address whether high-resolution orthorectified aerial imagery (i-e orthophotos) embeddings can substitute or complement high-quality, soil-specific variables to predict and explain multi-trophic soil biodiversity. First, we systematically evaluate the predictive performance of Light Gradient Boosting Machine (LGBM), Random Forest (RF), and Artificial Neural Networks (ANN) across four data configurations: (i) coarse-gridded environmental tabular data, (ii) high-quality in-situ tabular data, (iii) remote sensing-derived embeddings from EOF models, and (iv) a combination of embeddings and coarse-gridded environmental tabular data. This analysis aims to highlight whether image-derived embeddings alone or combined with gridded-environmental tabular data can match or surpass the predictive power of models relying on detailed in-situ soil-specific variables. Second, focusing on the best-performing model, we examine the efficiency of two EOF models (SatDINO and DOFA) to determine the added value of large-scale, self-supervised embeddings for soil biodiversity prediction, and explore the effect of dimensionality reduction through Principal Component Analysis (PCA). Third, we assess environmental drivers across data configurations and trophic groups in the French Alps, offering insight into how soil, climate, phenology, and landscape characteristics influence multi-trophic soil biodiversity.

## 2. Materials and Methods

### 2.1. Data Collection

#### 2.1.1. Soil Biodiversity Data

Soil trophic group abundance data, i.e., the number of DNA readings of the species found in the soil samples belonging to a trophic group, was obtained from the French Alps long-term observatory ORCHAMP [13] as detailed in [36] Soil samples were collected from 26 elevational gradients between 2016 and 2020. Each elevational gradient is characterized by uniform exposure and slope, includes plots of 30×30m, spaced 200m apart in altitude. Each plot was divided into 2x2m subplots, of which three random subplots were selected to extract composite soil samples (10 soil cores were collected and merged per subplot to form a composite sample). Soil samples were used for measuring the physicochemical properties of the soil and for eDNA extraction (15mg). DNA was extracted from soil using the methodology detailed in [22], followed by amplification with six DNA markers: two universal markers (euka02 for eukaryotes, bact01 for bacteria) and four clade-specific markers targeting fungi, insects, oligochaetes, and collembola. A bioinformatic pipeline was applied as described in [37] to process sequence data, remove contaminants, and assign molecular operational taxonomic units (MOTUs). Trophic and functional classifications were informed by databases such as FungalTraits, FUNGuild, and FAPROTAX, resulting in the assignment of MOTUs to 51 trophic groups across seven taxa: Bacteria, Fungi, Protist, Oligochaete, Insect, Collembola, and Metazoa. This process resulted in a dataset comprising 953 soil samples, each sample containing the respective abundance of each trophic group. A summary of descriptive abundance statistics is presented in Appendix A - Table A.2 for the 51 trophic groups.

#### 2.1.2. Environmental Covariates

In this study, we compiled a set of relevant environmental variables in tabular format from various sources, categorized into climate, soil, phenology, and landscape data (as detailed in Table 1).

**Table 1:** Summary of environmental covariates categorized by type (climate, soil, phenology, and landscape) and data source. The table lists each covariate along with its corresponding abbreviation, which is used throughout this study.

| Type | Source | Covariate | Abbreviation |
| --- | --- | --- | --- |
| Climate | CHELSA [38] | Mean annual air temperature | CLI.L.TMeanY |
|  |  | Temperature seasonality | CLI.L.TSeason |
|  |  | Annual precipitation amount | CLI.L.PTotY |
|  |  | Precipitation seasonality | CLI.L.PSeason |
|  |  | Total solar radiation | CLI.L.RTotY |
|  |  | Seasonal radiation | CLI.L.RSeason |
|  | SAFRAN-CROCUS [39] | Saturated water vapor pressure | CLI.H.Vpd |
|  |  | Rainfall rate | CLI.H.Rainf |
|  |  | Snowfall rate | CLI.H.Snowf |
|  |  | Nebulosity | CLI.H.Nebulosity |
|  |  | Surface incident direct shortwave radiation | CLI.H.DSWdown |
|  |  | Surface incident longwave radiation | CLI.H.LWdown |
|  |  | Surface incident diffuse shortwave radiation | CLI.H.SWdown |
|  |  | Relative humidity | CLI.H.Humrel |
|  |  | Reference crop evapotranspiration | CLI.H.ETP |
|  |  | Maximum daily temperature | CLI.H.TairMax |
|  |  | Minimum daily temperature | CLI.H.TairMin |
|  |  | Average daily temperature | CLI.H.TairMean |
|  |  | Wind | CLI.H.Wind |
|  |  | Temperature of the 1st cm of sun | CLI.H.Tg1 |
|  |  | Temperature about 10cm below the surface | CLI.H.Tg4 |
|  |  | Total snow depth | CLI.H.DTISBA |
| Soil | SOIL GRIDS [40] | Soil nitrogen | SOI.L.Nitro |
|  |  | Soil organic carbon | SOI.L.Soc |
|  |  | PH water | SOI.L.PhH2O |
|  |  | Clay | SOI.L.Clay |
|  |  | Silt | SOI.L.Silt |
|  |  | Sand | SOI.L.Sand |
|  | ORCHAMP [13] | Carbon to nitrogen ratio | SOI.H.cn |
|  |  | Soil moisture | SOI.H.Moist |
|  |  | Soil nitrogen | SOI.H.Nitro |
|  |  | Cation exchange capacity | SOI.H.Cec |
|  |  | Clay | SOI.H.Clay |
|  |  | Silt | SOI.H.Silt |
|  |  | Sand | SOI.H.Sand |
| Phenology | COPERNICUS Vegetation [41] | Seasonal productivity | PHE.Sprod |
|  |  | Seasonal length | PHE.Length |
|  |  | Seasonal amplitude | PHE.Ampl |
| Landscape | THEIA OSO Land Cover [42] | Broadleaved forests | LAN.BroadlForest |
|  |  | Conifer forests | LAN.ConiForest |
|  |  | Natural mineral surfaces | LAN.MineralSurf |
|  |  | Natural grasslands | LAN.Grass |
|  |  | Heathland | LAN.Shrub |
|  | BD TOPO® [43] | Water course | LAN.Water |

The climate data were sourced from CHELSA [38] and SAFRAN-CROCUS [39] datasets. CHELSA provides interpolated raster data with a spatial resolution of approximately 1 km, derived from downscaled ERA-Interim reanalysis outputs. We considered CHELSA as a coarse-gridded dataset. In contrast, SAFRAN-CROCUS offers higher-quality data obtained through reanalysis of in-situ meteorological station measurements, with a finer altitudinal resolution of approximately 300 meters. Similar than for climate, we derived soil variables from two sources that differed in terms of resolution and quality. First, we employed data from SOIL GRIDS [40] that provide global predictions of soil properties at a rather coarse resolution (1km) and moderate quality, and from ORCHAMP [13]. that provides high-quality in-situ measurements specific to our study area and characterized on the same samples than the eDNA data.

In addition to climate and soil variables, we incorporated phenology and landscape covariates in our study. Phenology variables were obtained from the Copernicus Vegetation [41] dataset, which provides daily vegetation indices at a 10-meter spatial resolution, derived from Sentinel-2 satellite observations. These variables include seasonal productivity, seasonal length, and seasonal amplitude, and are used to characterize the temporal dynamics of vegetation in the study areas. On the other hand, landscape variables were extracted from the THEIA OSO Land Cover [42] map and BD TOPO^®^ [43] from the Institut National de l’Information Géographique et Forestìere (IGN) of France. The first one provides land use classification maps for metropolitan France at a 10-meter spatial resolution, based on Sentinel-2 data. These variables include broadleaved forests, conifer forests, natural mineral surfaces, natural grasslands, and heathland. And the second one provides hydrographic features such as rivers, streams, and other flowing water bodies represented in our study by one variable (water course) that captures the proportion area around 100m of the plot crossed by a river/stream. All these variables offer detailed information on the composition and structure of the landscape in the study regions. They were considered to be of intermediate resolution and quality.

Finally, we incorporated orthophotographs from the BD ORTHO^®^ [44] dataset provided by the IGN. These images have a spatial resolution of 20 cm per pixel and we extracted them over the 26 elevational gradients studied, totaling 203 RGB images (Red, Green, and Blue bands) and 203 IRC images (Infrared, Red, and Green bands). The BD ORTHO^®^ dataset is a collection of high-resolution digital orthophotographs, rectified to ensure optimal geometric accuracy, and is updated every 3 to 4 years to reflect changes in the territory of France.

### 2.2. Experimental Setup

An overview of the methodology is illustrated in Fig. 1. The main objective is to predict the relative abundance of 51 trophic groups spanning 7 biological categories (Bacteria, Fungi, Protist, Oligochaete, Insect, Collembola, and Metazoa) across the French Alps. To achieve this, we developed independent regression models for each trophic group using three machine learning techniques (LGBM, RF, and ANN).

**Figure 1:**
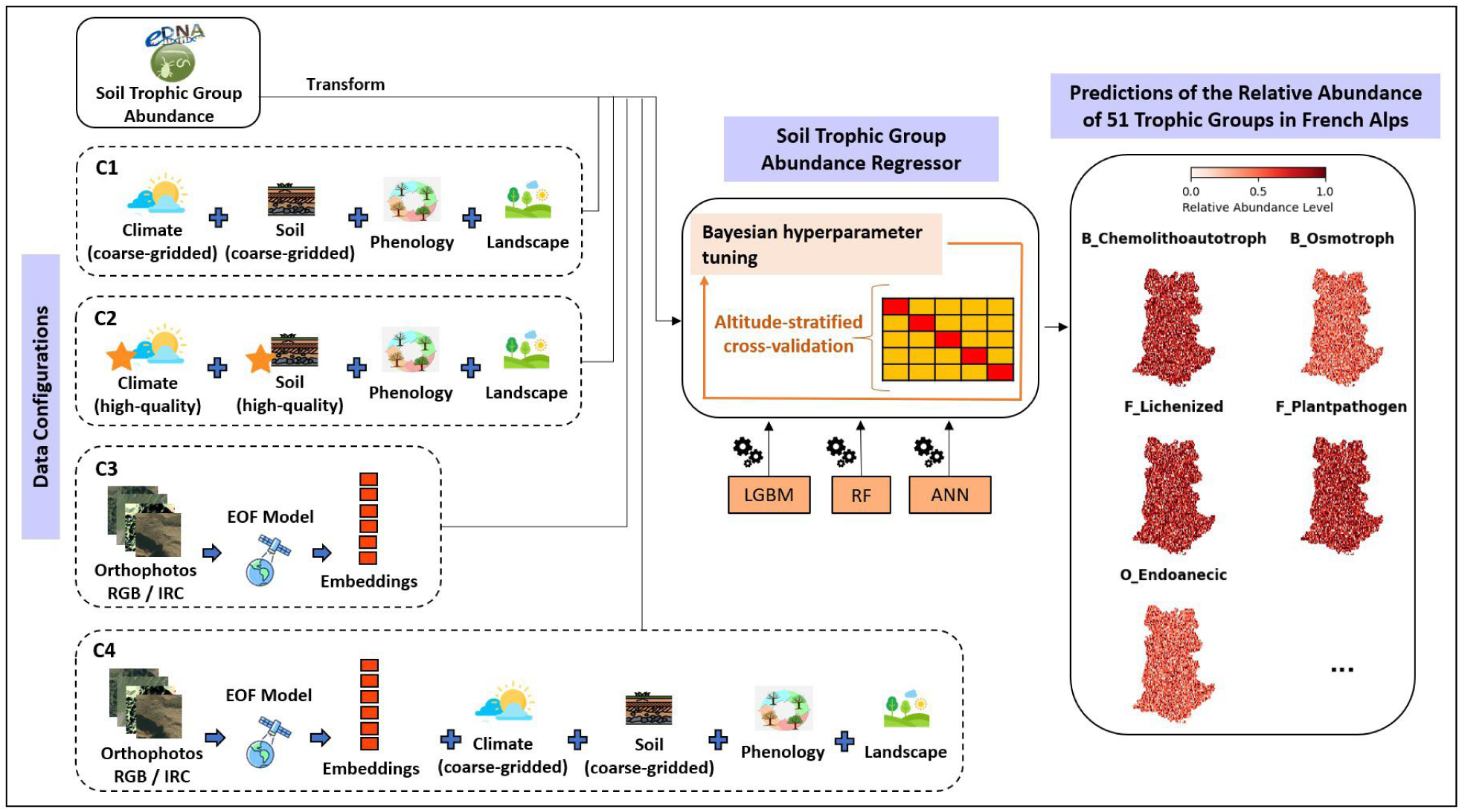
Overview of the methodology applied. The workflow consists of structuring environmental data into four configurations (C1–C4), applying machine learning models (LGBM, RF, ANN), and optimizing model performance through Bayesian hyperparameter tuning and altitude-stratified cross-validation. C1 and C2 are tabular datasets, where climate and soil variables are classified into coarse-gridded and high-quality in-situ data, while phenology and landscape variables provide complementary intermediate-scale information. C3 and C4 integrate remote sensing-derived embeddings from high-resolution orthophotographs, capturing fine-scale landscape features. The final predictions estimate the relative abundance of 51 trophic groups across the French Alps, supporting future biodiversity mapping efforts.

To evaluate the predictive power of different data sources and resolutions, we structured our input data into four configurations. Two of these configurations (C1 and C2) are purely tabular and differ in the resolution and source of climate and soil data, while the remaining two (C3 and C4) incorporate remote sensing-derived features.

Coarse-gridded tabular dataset (C1). This dataset integrates gridded climate and soil variables from CHELSA (∼1 km^2^ resolution) and SOIL GRIDS (∼250 m resolution), which provide large-scale, interpolated environmental predictions. Additionally, it includes phenology and landscape variables from COPERNICUS Vegetation, THEIA OSO Land Cover, and BD TOPO, which, despite their finer resolution ( 10m), originate from remote sensing and cartographic sources rather than direct field measurements.

- High-quality in-situ tabular dataset (C2). It incorporates higher-resolution, in-situ climate and soil variables from SAFRAN-CROCUS (∼300m resolution) and ORCHAMP (plot-level field measurements), which are derived from meteorological station data, reanalysis models, and direct soil sampling. To maintain consistency in landscape characterization, it includes the same phenology and land cover variables from COPERNICUS Vegetation, THEIA OSO Land Cover, and BD TOPO.
- Orthophoto-based embedding dataset (C3). It uses embeddings generated from pre-trained EOF models, applied to high-resolution BD ORTHO orthophotographs (20 cm spatial resolution). It captures fine-scale spatial patterns in the landscape, including vegetation structure and terrain features, which may influence soil biodiversity.
- Hybrid tabular-embedding dataset (C4). It combines coarse-gridded tabular data (C1) with embeddings extracted in C3.

By comparing C1 and C2, we aim to assess whether incorporating high-quality in-situ climate and soil data enhances the predictive accuracy of the models relative to coarse-gridded environmental data. The comparison between C2 and C3 evaluates the potential of orthophoto-based embeddings to capture landscape and environmental heterogeneity at a fine scale and determine whether they can serve as an alternative to high-quality in-situ predictors. Finally, the comparison between C2 and C4 examines whether integrating remote sensing-derived embeddings with coarse-gridded tabular data enhances model performance, potentially compensating for the absence of high-quality in-situ measurements.

#### 2.2.1. Generating Embeddings from Earth Observation Foundations Models

To incorporate high-resolution spatial information into our models, we employed two pre-trained EOF models to generate embeddings from high-resolution orthophotos of the BD ORTHO dataset:

- SatDINO [33]. It is a Vision Transformer (ViT) model specifically designed for high-resolution canopy height mapping, leveraging self-supervised learning. It incorporates the state-of-the-art DINOv2 training technique to produce robust, high-quality feature representations (embeddings) without the need for labeled data. The model was trained on RGB high-resolution satellite imagery from Maxar with a spatial resolution of 0.59 meters, covering geographically diverse regions like São Paulo (Brazil) and California (USA), i.e., diverse vegetation types, such as forests, mountainous terrains, and high degree of tree biodiversity. SatDINO is aligned with the mountainous focus of our study in the French Alps and is compatible with our orthophotos, which have a spatial resolution of 0.20 meters. We selected the Compressed SSLhuge checkpoint, which contains 749 million parameters and is optimized through quantization to reduce memory usage while enhancing computational efficiency. For input preparation, we used only the RGB orthophotos, cropping them to focus exclusively on the sampled plot regions. The images were adjusted to a resolution of 0.59 meters and resized to 224x224 pixels. SatDINO was configured to extract embeddings from the penultimate layer of the model, yielding an embedding size of 1280 for each input image.
- DOFA [32]. It is a ViT model designed to integrate multiple data modalities adaptively within a unified framework. The model incorporates training strategies such as masked image modeling, wavelength-conditioned dynamic patch embedding, and distillation-based multimodal continual pretraining. DOFA was trained on a diverse collection of datasets: Sentinel-1 (SAR, 2 channels), Sentinel-2 (multispectral, 9–13 channels), Gaofen (multispectral, 4 channels), NAIP (RGB, 3 channels), and EnMAP (hyper-spectral, 202 channels). This multimodal capability makes DOFA particularly suitable for applications requiring data fusion, such as ecological monitoring, land cover classification, and biodiversity studies.

DOFA aligns with our study due to its capability to process multimodal inputs. Given that our dataset includes infrared imagery (IRC images), we constructed 4-channel images (RGBI) by combining the RGB layers with the extracted infrared layer (I). These images were adjusted to match the resolution of 0.59 meters and resized to 224x224 pixels, ensuring compatibility with the model and facilitating direct comparison with SatDINO, which processes images under the same characteristics. For preprocessing, normalization was applied using the mean and standard deviation calculated from our image dataset, and wavelengths were defined based on standard RGBI values. We used the DOFA ViT base e100 checkpoint (111M parameters, non-quantized) to extract embeddings from the penultimate layer of the model. Each input image resulted in an embedding array of size 768, capturing spectral and spatial features for downstream analysis.

After generating the embeddings, they were integrated into our predictive models as predictors. These embeddings were used in cases C3 and C4. To evaluate the effect of different embedding sources, C3 was tested twice, once using only embeddings from SatDINO and once using only embeddings from DOFA. C4 extends C3 by combining the embeddings with the covariates from C1. Similar to C3, C4 was also evaluated twice, depending on whether the embeddings originated from SatDINO or DOFA.

This setup allows us to assess the predictive value of remote sensing embeddings, test their ability to replace tabular environmental variables, and determine whether their integration improves model performance.

#### 2.2.2. Data Preprocessing

For each taxa, we grouped its corresponding trophic groups and performed two normalization steps. The first normalization involved dividing the abundance of each trophic group within a sample by the total abundance of all trophic groups in the same sample (horizontally). This transformation converts absolute values into relative proportions, making the data comparable across samples. For the second normalization, we applied a min-max rescaling to each trophic group across all samples (vertically). This adjustment scaled the values proportionally between 0 and 1, ensuring that abundances are relative not only to the total within their sample but also to the observed range across all samples. Furthermore, we calculated two percentile thresholds (1% and 99%) for each trophic group. Values falling outside these thresholds were considered outliers and subsequently removed. The resulting values represent the relative abundances that we aim to predict. These steps mitigate magnitude biases in our models, facilitate convergence, better capture trends, and ensure the abundances are representative for each trophic group.

Since high-quality climate data are the only variables recorded daily, while the other covariates and abundance data are recorded annually, we computed the average of the high-quality climate variables over the last 10 years relative to the year in which abundance was measured. This ensures that all covariates are temporally aligned.

Finally, we concatenated all covariates, embeddings, and relative abundances. From this consolidated dataset, we organized the variables according to the four previously defined cases (C1, C2, C3, and C4), as illustrated in Fig. 1.

#### 2.2.3. Trophic Group Abundance Modeling

To model the four data configurations and identify the best model for each trophic group, we compared the performance of three machine learning techniques:

- Light Gradient Boosted Machine (LGBM) [45]. A gradient boosting decision tree algorithm that employs a leaf-wise tree growth strategy, significantly improving computational efficiency and reducing memory consumption. In our case, the hyperparameters tuned include ’max depth’ (1–20), ’n estimators’ (50–1000), ’learning rate’ (0.005–0.3), ’subsample’ (0.9–1), ’colsample bytree’ (0.5–1), and ’num leaves’ (20–200).
- Random Forest (RF) [46]. A bagging decision tree algorithm that constructs multiple decision trees and averages their predictions to enhance accuracy and reduce overfitting. The hyperparameters tuned for RF include ’max depth’ (1–20), ’n estimators’ (50–500), ’max features’ (0.2–1), and ’min samples leaf’ (1–10).
- Artificial Neural Network (ANN). A computational model with interconnected layers of neurons that learn patterns from data. The architecture consists of one MultiheadAttention layer, an intermediate dense layer with 256 neurons and ReLU activation, another dense layer with 512 neurons and a dropout rate of 0.01, and a final output layer with a single neuron. The model is trained for 1000 epochs using the AdamW optimizer with default weight decay. The loss function is SmoothL1Loss, which balances L1 and L2 loss components, making the model robust to outliers while smoothing small gradient variations. Additionally, we dynamically adjust the learning rate using ReduceLROnPlateau, which reduces the learning rate by a factor of 0.1 if loss minimization (mode=’min’) does not improve after 5 epochs (patience=5). The hyperparameters tuned for ANN include ’number of attention heads’ (2-8), ’learning rate’ (1e-7–1e-4), ’batch size’ (1–10), and ’early stopping threshold’ (5–30).

To navigate the hyperparameter space and determine the optimal configuration, we employed Hyperopt [47] with 50 evaluations. Hyperopt is a Bayesian optimization framework based on the Tree-structured Parzen Estimator (TPE) method. To guide the search, we used Mean Absolute Error (MAE) as the objective function, which ensures stability and prevents bias toward extreme predictions.

Before training the models, we stratified the samples to prevent data leakage and ensure a representative distribution of the samples in both training and test sets. Given that elevation strongly influences species distribution by affecting nutrient availability, temperature, humidity, and other environmental factors, stratification was performed in two steps. First, samples within each biological category were classified into five quantiles based on elevation. Second, we applied Stratified K-Fold cross-validation (K=5) using the pre-defined elevation-based classification and stored the sample indices for each training and testing set. This ensured that all techniques were evaluated using identical datasets.

The modeling process started by iterating over each taxon and then over each trophic group within that taxon. For each trophic group, we extracted the relevant data according to the configuration being tested (C1, C2, C3, or C4) and defined the hyperparameter search space using Hyperopt, based on the regression technique selected (LGBM, RF, or ANN). Then, we iterated through each stored fold, retrieving the training and testing indices to select the corresponding samples. For each fold, we extracted the target variable (relative abundance distribution of the trophic group) and computed its skewness coefficient. If the skewness exceeded 1, we applied a logarithmic transformation to reduce asymmetry and improve model stability; otherwise, we left the data unchanged. We also preprocessed the predictor variables as follows: tabular covariates (climate, soil, phenology, and landscape) were normalized using min-max transformation and embeddings remained unchanged. The models were trained using the preprocessed data and hyperparameters selected by Hyperopt. After obtaining predictions, we reversed the logarithmic transformation applied to the target variable. We stored predictions from all folds and evaluate model performance using four metrics: Root Mean Square Error (RMSE) to penalize large errors; MAE to assess the average prediction error; Coefficient of Determination (R² score) to measure the explanatory power of the models; and Spearman’s Rank Correlation Coefficient (*ρ*) to capture the monotonic relationship between predictions and true values. This process was repeated iteratively with new hyperparameter configurations until MAE is minimized, allowing us to identify the best-performing model for each trophic group.

By systematically comparing the four data configurations, we quantified the relative contributions of tabular environmental covariates and high-resolution embeddings in predicting trophic group abundance, providing insights into the added value of remote sensing-derived features.

## 3. Results

### 3.1. Comparative Performance of Machine Learning Techniques for Different Data Configurations

To identify the best approach to predict trophic group abundance, we evaluated the performance of three different ML techniques (RF, LGBM, and ANN) and four data configurations (C1, C2, C3, and C4), where cases C3 and C4 have been tested using embeddings generated by the SatDINO model. RF consistently achieved the highest predictive performance, slightly outperforming LGBM (Fig. 2, see Appendix B for R², MAE, and RMSE results). However, LGBM remained computationally more efficient, executing significantly faster than both RF and ANN across all data configurations.

**Figure 2:**
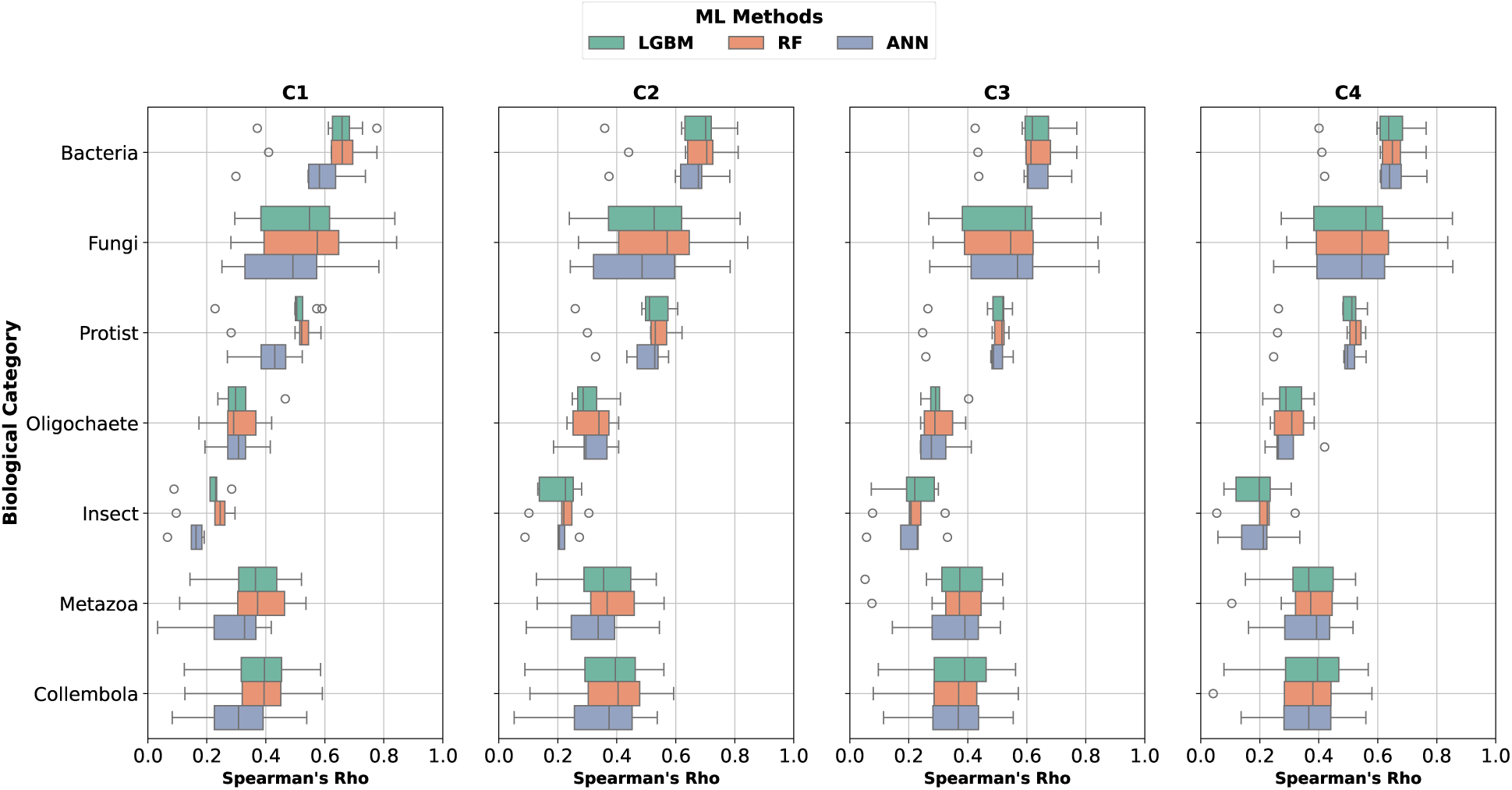
Comparison of model performance across trophic groups using Spearman’s correlation. The boxplots summarize the predictive performance for each biological category (Bacteria, Fungi, Protist, Oligochaete, Insect, Metazoa, and Collembola) across the four data configurations (C1–C4). Each configuration was evaluated using three machine learning models (LGBM, RF, and ANN). Higher values indicate stronger agreement between predicted and observed relative abundance.

Beyond differences in model architecture and efficiency, we also observed notable variation in predictive performance across biological categories. More precisely, Bacteria, Fungi, and Protists exhibited the highest predictability, as indicated by consistently higher Spearman’s correlation values (Fig. 2). These groups likely benefited from stronger associations with environmental variables due to their high abundance and relatively well-defined ecological niches. Metazoa and Collembola showed intermediate predictive performance, suggesting a more variable response to the environmental drivers considered. In contrast, Oligochaete and Insect groups showed the lowest performance.

Regarding data configurations, we found that the overall mean value of models trained on high-quality in-situ tabular data (C2) using RF outperformed all other configurations (see Appendix B, Fig. B.12), confirming that high-resolution climate and soil variables significantly improve predictive accuracy. While models only based on embedding data (C3) captured relevant landscape information, they failed to surpass models based on C1, C2, or C4.

Regarding specific trophic groups variability, in most of the groups within Metazoa and Insects, where tabular data alone struggles, C4 resulted in better predictions (Fig. B.12). In these cases, the inclusion of embeddings in C4 improved predictive performance relative to tabular-only models, indicating that the additional image-derived features contributed complementary information beyond the tabular covariates. Moreover, not all trophic groups within a biological category perform equally well, with some groups showing striking disparities in predictability. For instance, in Bacteria, models for B Zooparasite achieved Spearman correlation values consistently above 0.751 across all models nearly twice as high as B Osmotroph, where models struggled below 0.437. A similar pattern was observed for Fungi, where models for F Ectomycorrhizal reached values above 0.783, whereas models for F Lichenparasite performed nearly three times worse, with values plummeting below 0.292. The gap was even more pronounced in Metazoa, where models for M NHerbivores reached values above 0.50, while models for M NAnimalParasites dropped more than threefold to a mere 0.162. In Collembola, models for C Euedaphic reached 0.537, whereas predictions for C Hemiedaphic lagged nearly four times behind, struggling at just 0.137.

### 3.2. Relevance of Orthophoto Embeddings to predict Soil Biodiversity

Since models based on C3 underperformed compared to tabular-based approaches in most cases, we further investigated whether either the choice of embedding model or the application of dimensionality reduction could enhance their predictive utility. To this end, we compared predictive performance across embedding-based data configurations (C3, C4), this time using embeddings derived from both SatDINO and DOFA. Both configurations were evaluated with and without dimensionality reduction through PCA (C3+PCA, C4+PCA). To implement PCA, we extended the training pipeline described in 2.2.3 by treating the number of components as an additional hyperparameter. Embeddings were standardized to zero mean and unit variance and projected into a reduced-dimensional space accordingly, without altering the overall cross-validation or model optimization procedure. Results indicated that no embedding-based model (C3, C4) surpassed the best tabular-based model (C2) in predictive performance Fig. 3. Although embeddings contributed relevant spatial information, they did not provide sufficient complementary features to outperform high-quality in-situ tabular data. Among embedding-based approaches, in most cases models using SatDINO-derived embeddings outperformed slightly those using DOFA-derived embeddings, suggesting that its learned representations better capture environmental patterns relevant to soil biodiversity. Given this trend, subsequent analyses in this study focus on SatDINO embeddings.

**Figure 3:**
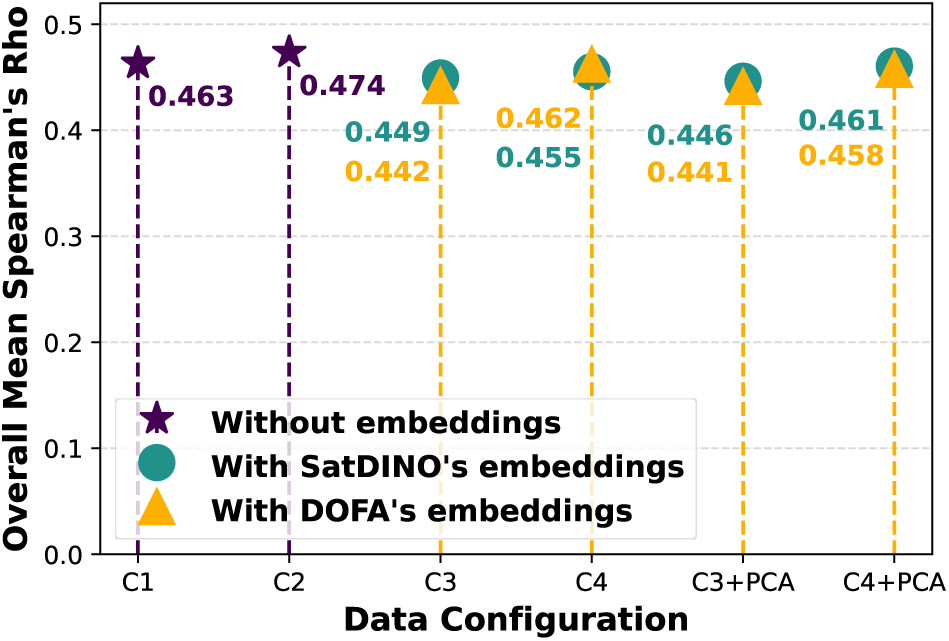
Comparison of overall mean Spearman’s correlation for tabular-based and embedding-based models using Random Forest. C1 and C2 represent the best-performing tabular models, while C3 and C4 incorporate embeddings from SatDINO and DOFA, with and without dimensionality reduction via PCA (C3+PCA, C4+PCA).

To further understand the extent to which embeddings captured meaningful ecological patterns that contribute to predictive performance, we examined how they encoded landscape information. A PCA projection of SatDINO embeddings (Fig. 4) illustrates the degree to which they differentiate among dominant land cover types. The observed projection suggested that SatDINO embeddings partially differentiate between land cover types, particularly for Coniferous Forests and Mineral Surfaces, while Grasslands and Shrubs exhibit greater overlap, likely due to similar spectral and structural characteristics.

**Figure 4:**
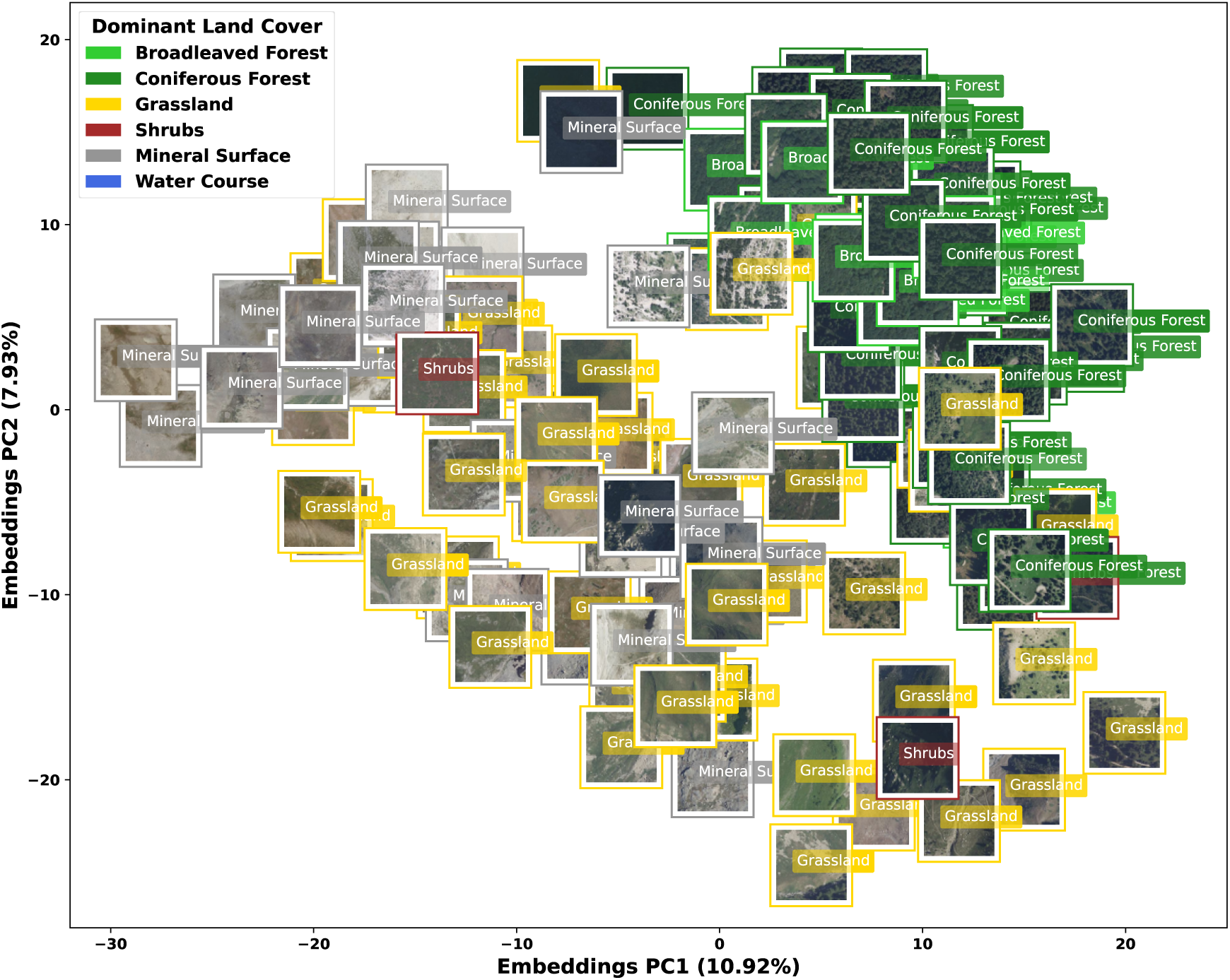
PCA projection of SatDINO embeddings with land cover visualization. Each point represents a sample in the reduced-dimensional embedding space, overlaid with its corresponding RGB orthophoto and framed according to the dominant land cover type. The figure illustrate the extent to which SatDINO embeddings capture landscape variability.

Furthermore, we observed a substantial alignment between high-quality tabular data (C2) and SatDINO embeddings, as revealed by the continuity of gradient patterns in the PCA projection colored by the first embedding component (Fig. 5, left). This suggests that major spatial structures captured by the embeddings are already encoded in the tabular variables. However, the second embedding component (Fig. 5, right) exhibited weaker gradient structure, indicating that some additional, albeit limited, variability may be uniquely encoded in the embeddings. Overall, this partial redundancy explains why incorporating embeddings into models trained on coarse-gridded data (C4) did not improve predictive performance.

**Figure 5:**
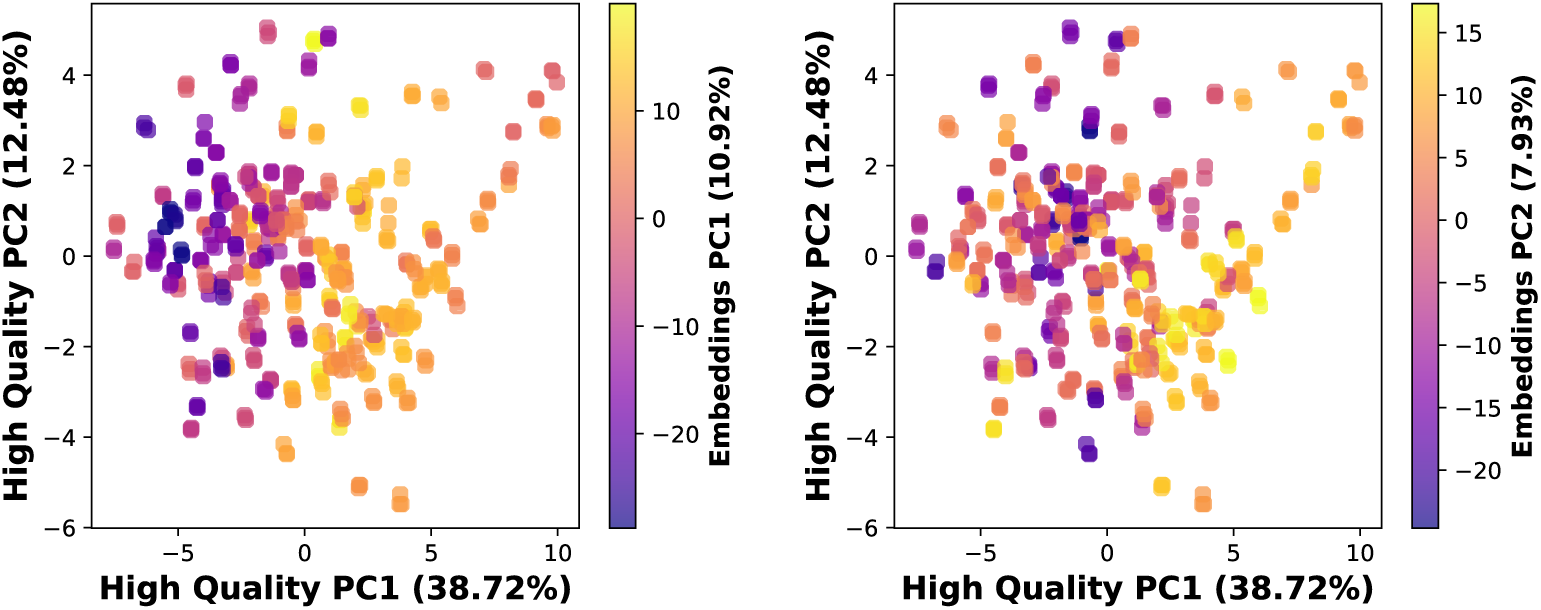
Relationship between high-quality tabular data and SatDINO embeddings through PCA projections. Both plots display the first two principal components of the high-quality in-situ tabular dataset (C2 configuration). The left plot is colored by the first principal component of SatDINO embeddings (PC1), while the right one by the second principal component of SatDINO embeddings (PC2). The plots illustrate the extent to which patterns in high-quality tabular data correspond to variations in embedding space.

Together, these results indicated that while SatDINO-derived embeddings captured landscape variability, they did not introduce sufficiently novel information beyond what was already encoded in high-resolution climate and soil variables.

### 3.3. Evaluating Environmental Drivers and Spatial Patterns in Soil Biodiversity Across Data Configurations

The relative importance of environmental predictors varied across data configurations and trophic groups (Fig. 6). In most cases, climate and soil variables exhibited the highest relative importance in C1 (coarse-gridded tabular data) and C2 (high-quality in-situ tabular data), as seen in Fig. 6. This aligns with our PCA analyses based on coarse-gridded tabular data (C1) and high-quality in-situ data (C2), respectively (Fig. 7), where climate and soil variables accounted for the largest proportion of variance, confirming their dominant role in structuring soil biodiversity. Additionally, when comparing covariates from C1 and C2 (Fig. 7), both datasets revealed a clear altitudinal gradient, with samples clustering along the first axis according to their elevation (yellow/green points appearing more separated). This correlation was strongest for high-quality insitu data, which explained a greater proportion of variability (51.2%), indicating that finer-resolution climate and soil variables better capture key ecological gradients. Notably, predictor contributions differed between configurations: SOI.L.Nitro and SOI.L.PhH2O were strong contributors in C1 but showed reduced importance in C2, while high-resolution temperature extremes (CLI.H.TairMax, CLI.H.TairMin) gained prominence in C2 7, reinforcing their critical role in shaping soil communities. In contrast, when incorporating embeddings in C4 (hybrid tabular-embedding dataset), climate variables largely retained their dominant role, while the relative importance of soil features declined as embeddings gained influence (Fig. 6). This shift occurs because embeddings capture certain spatial or structural aspects of the environment that partially overlap with soil predictors. Notably, P Photoautotroph exhibited the highest reliance on orthophoto embeddings, whereas the best-performing model for F Ectomycorrhizal assigned minimal importance to them while achieving the highest predictive accuracy.

**Figure 6:**
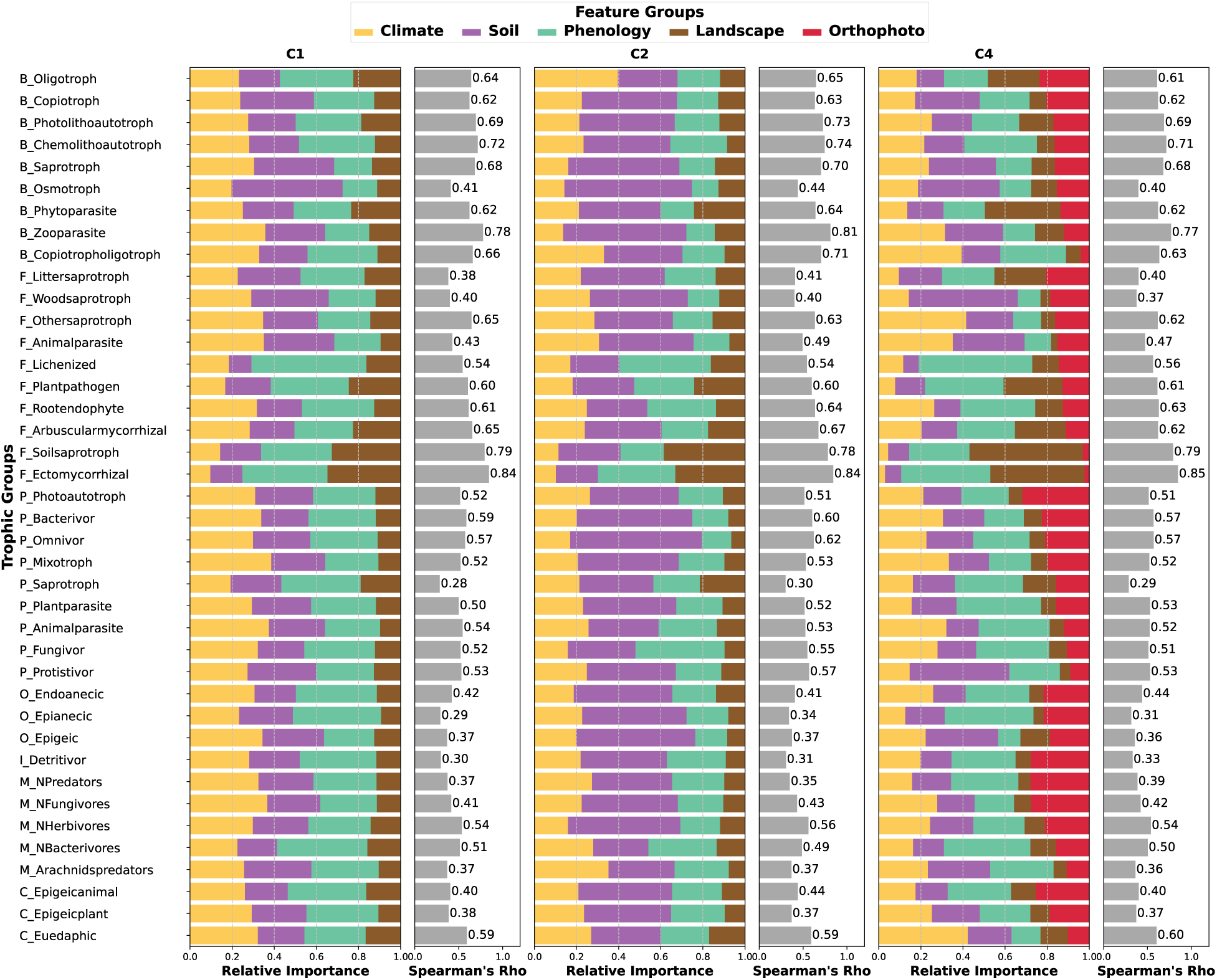
Comparison of feature importance across data configurations (C1, C2, C4) for trophic groups with models that achieved Spearman’s correlation *>* 0.3 in C2. The left column in each configuration represents the relative importance of feature groups (climate, soil, phenology, and embeddings), while the right column displays model performance obtained with Random Forest. Feature importance values are averaged within each group, and trophic groups are sorted by the contribution of embeddings (red bars) from highest to lowest. In C4, embeddings were derived from SatDINO and reduced via PCA. This visualization highlights feature relevance shifts across data configurations.

**Figure 7:**
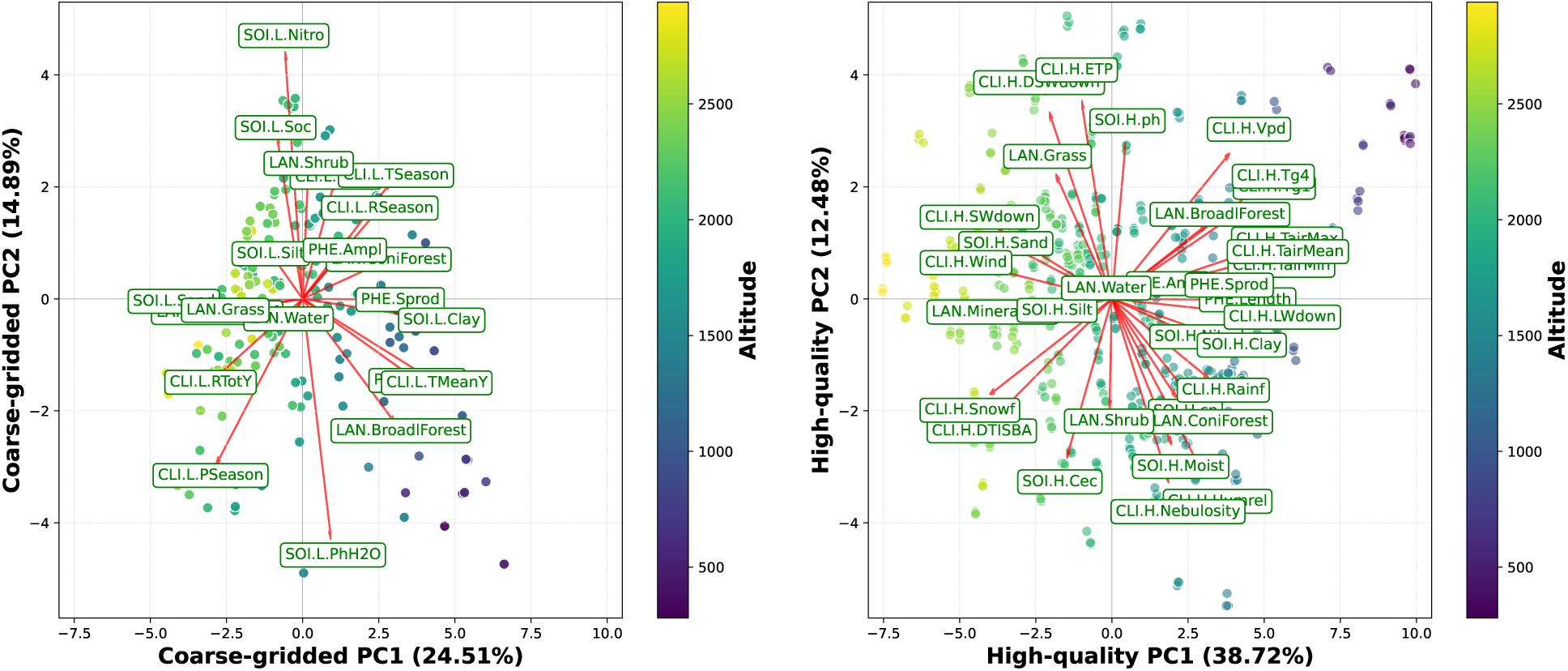
Biplots of environmental variable contributions based on in C1 (coarse-gridded tabular data, left) and C2 (high-quality in-situ tabular data, right). Each point represents a sample projected onto the first two principal components, with colors indicating elevation.

To further explore the relationship between environmental structure and trophic group distributions, we examined spatial patterns of predicted abundance for F Ectomycorrhizal and B Zooparasite (Fig. 8). For F Ectomycorrhizal, both embedding-based and tabular models exhibited a well-defined spatial gradient, with higher predicted abundances in Coniferous Forests and lower values in Grasslands (Fig. 8). This pattern aligned with the ecological niche of ectomycorrhizal fungi, which form mutualistic associations with tree roots and are primarily found in forested environments. The strong agreement between only embedding-based and tabular models in predicting the dominant land cover types suggests that embeddings can, in some cases, capture key habitat associations. Conversely, B Zooparasite predictions lacked spatial organization in the embedding-based model, showing no clear gradient and weak clustering (Fig. 8). Unlike ectomycorrhizal fungi, zooparasitic bacteria may be more influenced by host species distribution, microhabitat conditions, or other biotic interactions that are not well captured by embeddings. Additionally, the discrepancy in land cover associations between embedding-based and tabular models suggests that embeddings not effectively encode the ecological drivers of this trophic group.

**Figure 8:**
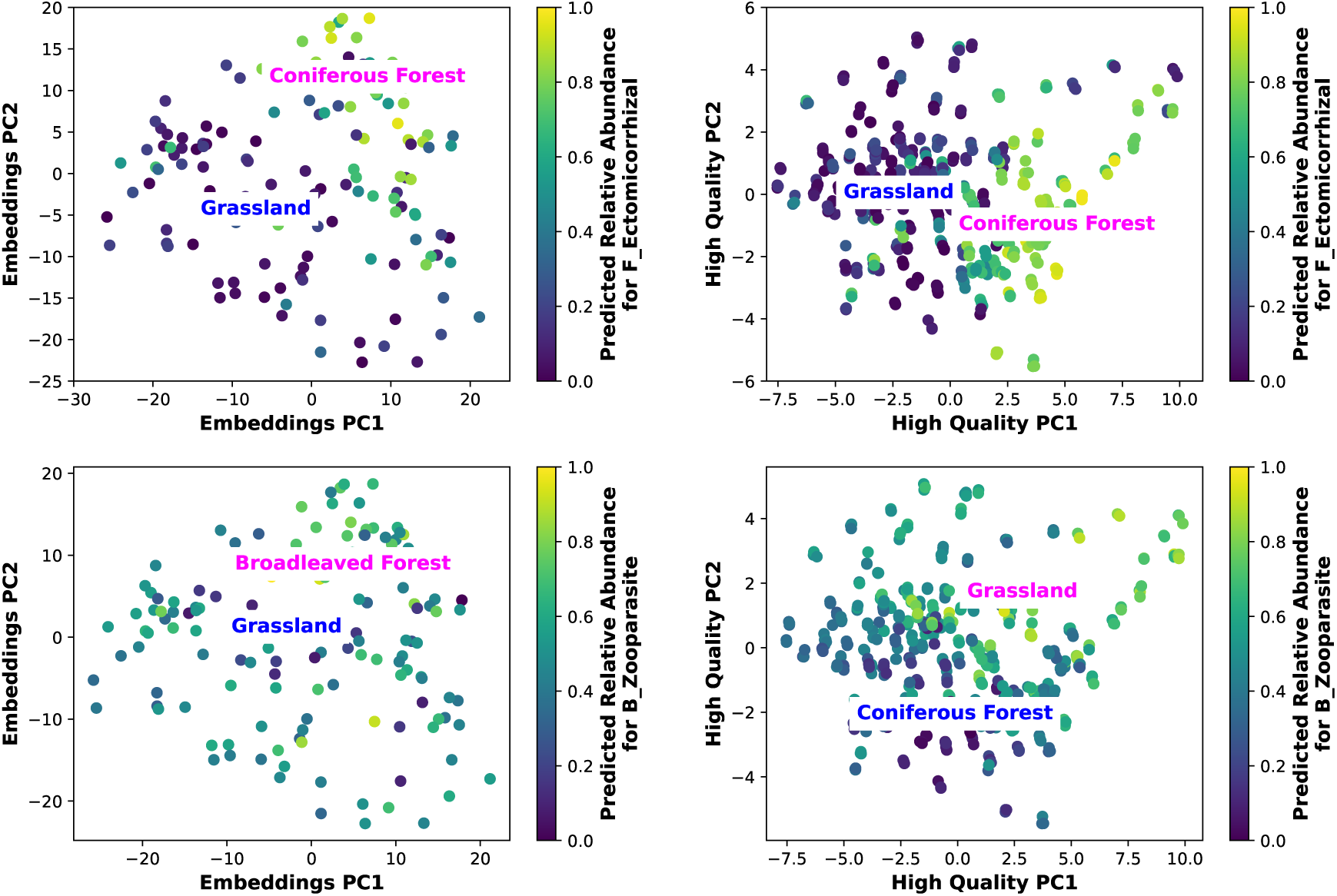
Spatial patterns in predicted relative abundance for *F Ectomycorrhizal* (top) and *B Zooparasite* (bottom). Each pair of plots compares the best predictive models using SatDINO embeddings (left - configuration C3) and high-quality tabular data (right - configuration C1). To visualize spatial patterns, embeddings and tabular data were reduced via PCA, and the first two principal components were plotted. Color gradients represent predicted relative abundance, while blue and magenta labels indicate the dominant land cover types associated with the lowest and highest predicted abundances, respectively.

## 4. Discussion

This study significantly advances our understanding of predicting soil biodiversity by comparing highquality in-situ environmental measurements, coarse-gridded predictors, and remote sensing-derived embeddings through advanced machine learning techniques. Our key finding is that high-resolution, in-situ climate and soil variables remain superior predictors of soil biodiversity, emphasizing their critical role in accurate ecological modeling. Nevertheless, remote sensing-derived embeddings can complement these predictors, especially where direct in-situ measurements are scarce.

Our results showed clear variation in predictive accuracy across the seven biological categories evaluated (2). Microbial groups such as Bacteria, Fungi, and Protists exhibited consistently higher predictive performance, reflecting their stronger, more direct relationships with soil and climate conditions. In contrast, Metazoa and Collembola presented intermediate predictability, while Insect and Oligochaete groups proved challenging due to greater ecological complexity, broader habitat use, and lower data availability. For instance, ectomycorrhizal fungi were particularly well-predicted (B.12). These fungi form obligate mutualistic relationships with tree roots, primarily in temperate and boreal forest ecosystems, where their presence and abundance are closely tied to host tree distribution. Since forested environments are well characterized by climate, soil, and landscape variables, models could effectively capture the environmental conditions that drive ectomycorrhizal abundance. Conversely, hemiedaphic Collembola (C Hemiedaphic microorganismconsumers) showed the lowest predictability (B.12). Hemiedaphic species exhibit greater vertical mobility, transitioning between the soil surface and deeper layers in response to microclimatic conditions. This dynamic habitat use makes their abundance highly variable and context dependent, potentially leading to weaker associations with coarse scale environmental predictors. This aspect was reflected in our results, where models relying on coarse-gridded tabular data (C1) and even high-resolution remote sensing-derived embeddings (C3 and C4) did not improve upon those trained with high-quality in-situ tabular data (C2) (3). While the raw imagery offers a resolution of 20cm, the learned embeddings may not fully retain or translate this spatial detail into ecologically meaningful features for belowground taxa (3). These findings highlight the critical role of not only spatial resolution but also ecological relevance in feature representation, suggesting that coarse-gridded or non-specific representations may obscure key environmental drivers. Nevertheless, embeddings were not without value; in some cases, they captured spatial aspects of the environment that partially overlapped with soil-related predictors(4 and 5). For instance, F Ectomycorrhizal assigned minimal importance for embeddings where as P Photoautotroph showed the highest reliance on orthophoto-derived embeddings (6). This discrepancy highlights group-specific relevance of embeddings, where some trophic groups may benefit from remote sensing-derived features, while others remain more dependent on direct soil and climate measurements, i.e., model architectures and feature representations may need to be tailored to the ecological traits of each functional group. Additionally, our models treat each trophic group independently, overlooking potential ecological interactions such as trophic dependencies, which could refine abundance predictions. Accounting for trophic dependencies might indeed help predicting secondary consumers or predators [14]). Second, environmental variables overlook biological processes like competition, symbiosis, and dispersal, yet integrating ecological networks or meta-community models could enhance predictive accuracy.

The strong predictive power of in-situ environmental variables aligns with previous studies highlighting the necessity of fine-scale soil measurements to model biodiversity, as soil properties exhibit high spatial heterogeneity that shapes species distributions and ecological processes at local scales. Variables such as soil pH, texture, organic matter content, and moisture availability play fundamental roles in structuring soil communities and demand fine spatial resolution to be effectively captured. [48]. Additionally, SDMs improve their predictive performance when high-resolution predictors are used, particularly for sessile organisms such as soil microbes and plants and, in contrast, coarser-resolution environmental predictors can lead to significant misrepresentation of biodiversity patterns [49]. Among the key soil variables that require high spatial resolution for biodiversity modeling and mapping, soil pH emerges as a dominant factor influencing microbial community distributions, as demonstrated in [23]. Other critical variables include soil texture (clay, silt, and sand fractions), organic carbon content, and soil moisture availability, all of which play fundamental roles in structuring soil biological communities [50, 36].

In contrast, there are studies where high-resolution satellite-derived variables improve broad-scale biodiversity mapping [51, 52], as they provide continuous and standardized measurements of vegetation cover, land use, and climatic conditions at a global scale. These datasets have proven useful for identifying largescale biodiversity patterns and monitoring ecosystem changes over time [53]. However, when attempting to predict abundance at finer spatial scales, particularly for soil communities, we observed a lack of specificity in capturing the heterogeneity of belowground environments. This is demonstrated in 4, where the embedding space showed limited separation between land cover types, especially those with similar spectral or structural characteristics. This is because soil biodiversity is strongly influenced by micro-scale variations in soil chemistry, moisture, and organic matter—factors that satellite-derived features struggle to resolve with the necessary precision [54, 55]. While remote sensing can infer proxies for soil conditions through vegetation indices and land surface temperature, it lacks direct measurements of crucial edaphic properties, leading to reduced predictive power. Consequently, relying exclusively on remote-sensing data imposes limitations that fail to capture the complexity of soil biodiversity drivers. This reinforces the observation that, although remote sensing-derived embeddings provide relevant spatial features, they cannot fully replace high-quality in-situ tabular environmental predictors in soil biodiversity modeling.

Finally, this limitation is reflected in the performance of embeddings derived from both remote sensing models evaluated, SatDINO and DOFA. Despite their architectural and methodological differences, embeddings from both models yielded similar predictive outcomes, with only a slight advantage for SatDINO. This raises the question of whether embeddings from different models truly capture distinct environmental information or if they converge toward a shared representation of landscape features. One potential explanation for the limited predictive utility of embeddings, extracted from SatDINO and DOFA, lies in how they encode environmental heterogeneity. Studies in representation learning [56, 57] suggest that while embeddings trained on different data modalities may initially capture distinct information, they often converge toward a shared statistical representation of reality. This means that despite their differences in architecture and input modalities, both SatDINO and DOFA may ultimately extract highly similar landscape features, limiting the diversity of information available for predictive modeling. As a result, their embeddings might not introduce novel environmental variability beyond what is already represented in high-quality in-situ tabular data.

## 5. Conclusion and Perspectives

Our study demonstrates that the predictive performance of machine learning models for soil biodiversity varies significantly across trophic groups, reflecting both taxon-specific ecological complexity and methodological challenges. Among the tested approaches, models trained exclusively with tabular environmental data, particularly those incorporating high-quality in-situ measurements, consistently achieved the strongest predictive performance. In contrast, while high-resolution orthophoto embeddings derived from foundation models captured spatial structure, they did not fully reproduce the fine-scale edaphic signals required to predict belowground biodiversity. This suggests that the current generation of image-based embeddings, despite their sub-meter input resolution, may not yet encode ecologically relevant features with sufficient fidelity for all soil taxa. Nonetheless, while orthophoto embeddings did not outperform tabular environmental covariates, they provided a viable alternative in scenarios where traditional field-based data collection is infeasible or limited. These findings emphasize the need for a dual strategy that leverages either comprehensive tabular environmental data or remote sensing-derived embeddings, depending on data availability and the specific constraints of a given study. Such an approach enhances the flexibility and scalability of biodiversity predictions, particularly in regions where high-resolution environmental datasets are scarce.

To further improve soil biodiversity modeling, future research could explore several key directions. One is the incorporation of multispectral data from Sentinel-2 and other Earth Observation satellites, which could improve the characterization of environmental conditions, particularly for large-scale biodiversity mapping. Additionally, developing predictive models that explicitly leverage trophic network data could capture source-target relationships, enhancing understanding of interdependencies among soil organisms and potentially enhancing predictive accuracy. Furthermore, integrating ecological networks and metacommunity models could help account for species interactions such as competition, symbiosis, and dispersal, which are often overlooked in purely environmental approaches. Finally, experimenting with transformer-based architectures and other deep learning frameworks designed for multimodal data fusion could enable better integration of tabular, image, and network-based information, capturing the complexity of soil biodiversity patterns more effectively.

## Acknowledgments

This work was co-funded by the European Union (Natura Connect, No: 101060429 and OBSGESSION, No.: 101134954) and the Agence Nationale pour la Recherche through the MIAI@Grenoble Alpes (ANR-19-P3IA-0003) institute and the Office Fraņcais de la Biodiversité (OFB). Content reflects only the views of the project owners

## Appendices

Additional details omitted from the main text are provided here as supplementary material.

## Appendix A. Summary Statistics of Trophic Group Abundances

This appendix provides summary statistics of trophic group abundances used for modeling.

**Table A.2:** Summary of eDNA read counts for 51 trophic groups across 953 soil samples collected in the French Alps. Trophic groups are organized within seven biological categories: Bacteria, Fungi, Protists, Oligochaetes, Insects, Metazoa, and Collembola. The table reports the mean read count (Avg.), standard deviation (Std.), and observed range (Min.-Max.) for each group.

| Bio. Categ. | Trophic Group | Avg. | Std. | Min. | Max. |
| --- | --- | --- | --- | --- | --- |
| Bacteria | B_chemolithoautotroph | 56.25 | 60.34 | 0 | 769 |
|  | B_copiotroph | 1757.11 | 605.95 | 60 | 5408 |
|  | B_copiotropholigotroph | 318.43 | 136.17 | 0 | 810 |
|  | B_oligotroph | 188.16 | 177.60 | 0 | 1519 |
|  | B_osmotroph | 7.97 | 33.54 | 0 | 466 |
|  | B_photolithoautotroph | 9.63 | 18.99 | 0 | 480 |
|  | B_phytoparasite | 298.02 | 190.01 | 0 | 1779 |
|  | B_saprotroph | 663.33 | 283.38 | 1 | 1517 |
|  | B_zooparasite | 281.05 | 207.46 | 0 | 3592 |
| Fungi | F_animalparasite | 51.26 | 90.36 | 0 | 830 |
|  | F_arbuscularmycorrhizal | 1400.33 | 2710.40 | 0 | 15114 |
|  | F_ectomycorrhizal | 3626.58 | 4094.26 | 0 | 14111 |
|  | F_lichenized | 19.15 | 121.26 | 0 | 2108 |
|  | F_lichenparasite | 20.87 | 101.08 | 0 | 2552 |
|  | F_littersaprotroph | 970.39 | 1302.17 | 0 | 12010 |
|  | F_mycoparasite | 23.89 | 47.68 | 0 | 607 |
|  | F_othersaprotroph | 243.88 | 460.36 | 0 | 4745 |
|  | F_plantpathogen | 99.19 | 139.20 | 0 | 936 |
|  | F_rootendophyte | 49.57 | 181.13 | 0 | 3300 |
|  | F_soilsaprotroph | 3905.71 | 3662.94 | 0 | 14708 |
|  | F_woodsaprotroph | 476.12 | 888.69 | 0 | 8476 |
| Protist | P_animalparasite | 8.83 | 15.04 | 0 | 181 |
|  | P_bacterivor | 83.61 | 40.16 | 0 | 200 |
|  | P_fungivor | 3.60 | 4.72 | 0 | 29 |
|  | P_mixotroph | 60.43 | 29.04 | 0 | 252 |
|  | P_omnivor | 142.82 | 57.42 | 0 | 378 |
|  | P_photoautotroph | 14.94 | 13.30 | 0 | 136 |
|  | P_plantparasite | 6.70 | 13.06 | 0 | 186 |
|  | P_protistivor | 15.84 | 11.92 | 0 | 82 |
|  | P_saprotroph | 0.54 | 12.37 | 0 | 378 |
| Oligochaete | O_Endoanecic | 729.69 | 2732.25 | 0 | 18467 |
|  | O_Endogeic | 165.19 | 1261.93 | 0 | 18210 |
|  | O_Epianecic | 623.89 | 2488.93 | 0 | 18472 |
|  | O_Epigeic | 515.03 | 2055.58 | 0 | 18049 |
|  | O_Intermediate | 644.81 | 2543.35 | 0 | 18472 |
| Insect | I_detritivor | 23.17 | 25.97 | 0 | 66 |
|  | I_fungivor | 0.79 | 5.67 | 0 | 66 |
|  | I_omnivor | 1.30 | 8.06 | 0 | 66 |
|  | I_phytophage | 12.82 | 21.63 | 0 | 66 |
|  | I_predator | 9.71 | 19.37 | 0 | 66 |
| Metazoa | M_Arachnidspredators | 23.03 | 78.26 | 0 | 1010 |
|  | M_NAnimalparasites | 0.92 | 15.70 | 0 | 469 |
|  | M_NBacterivores | 61.67 | 100.31 | 0 | 1273 |
|  | M_NFungivores | 11.08 | 21.67 | 0 | 356 |
|  | M_NHerbivores | 12.15 | 31.48 | 0 | 732.50 |
|  | M_NOmnivores | 1.73 | 11.12 | 0 | 221 |
|  | M_NPredators | 3.16 | 9.05 | 0 | 139 |
| Collembola | C_Epigeicanimal_andmicroorganismsconsumers | 81.90 | 159.92 | 0 | 688 |
|  | C_Epigeicplant_andmicroorganismconsumers | 149.27 | 205.73 | 0 | 688 |
|  | C_Euedaphic_microorganismconsumers | 230.76 | 239.22 | 0 | 688 |
|  | C_Hemiedaphic_microorganismconsumers | 0.76 | 7.63 | 0 | 142 |

## Appendix B. Comparative Performance of ML Techniques and Data Configurations

The following figures present additional evaluation metrics such as R² score, MAE, and RMSE, for each ML technique (RF, LGBM, ANN) across the four data configurations (C1 to C4). Also, we present a heatmap of Spearman correlations across all trophic groups, techniques, and data configurations. This figure provides a more granular view of model performance.

**Figure B.9:**
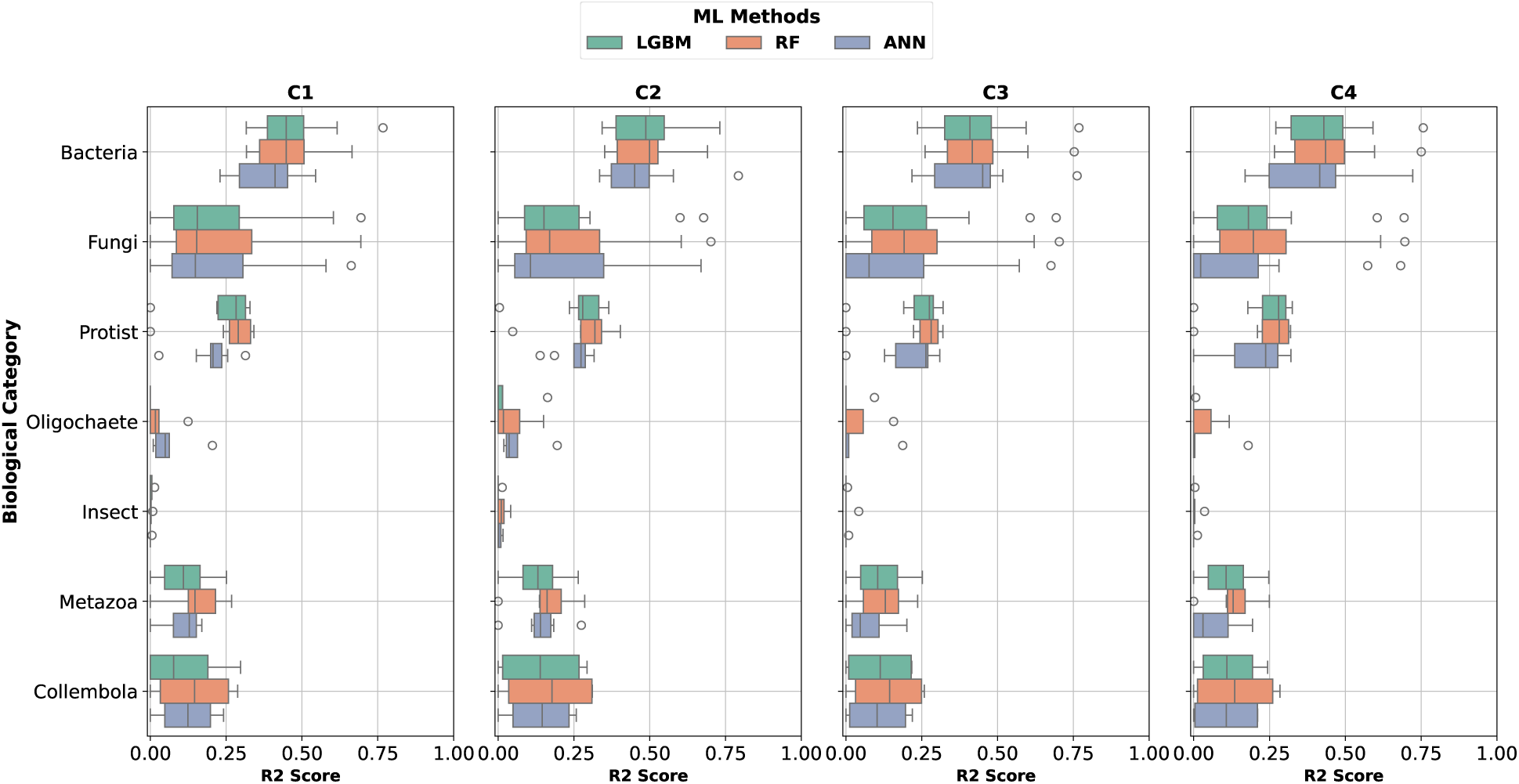
Comparison of model performance across trophic groups using R² score. The boxplots summarize the predictive performance for each taxon (Bacteria, Fungi, Protist, Oligochaete, Insect, Metazoa, and Collembola) across the four data configurations (C1–C4). Each configuration was evaluated using three machine learning models (LGBM, RF, and ANN).

**Figure B.10:**
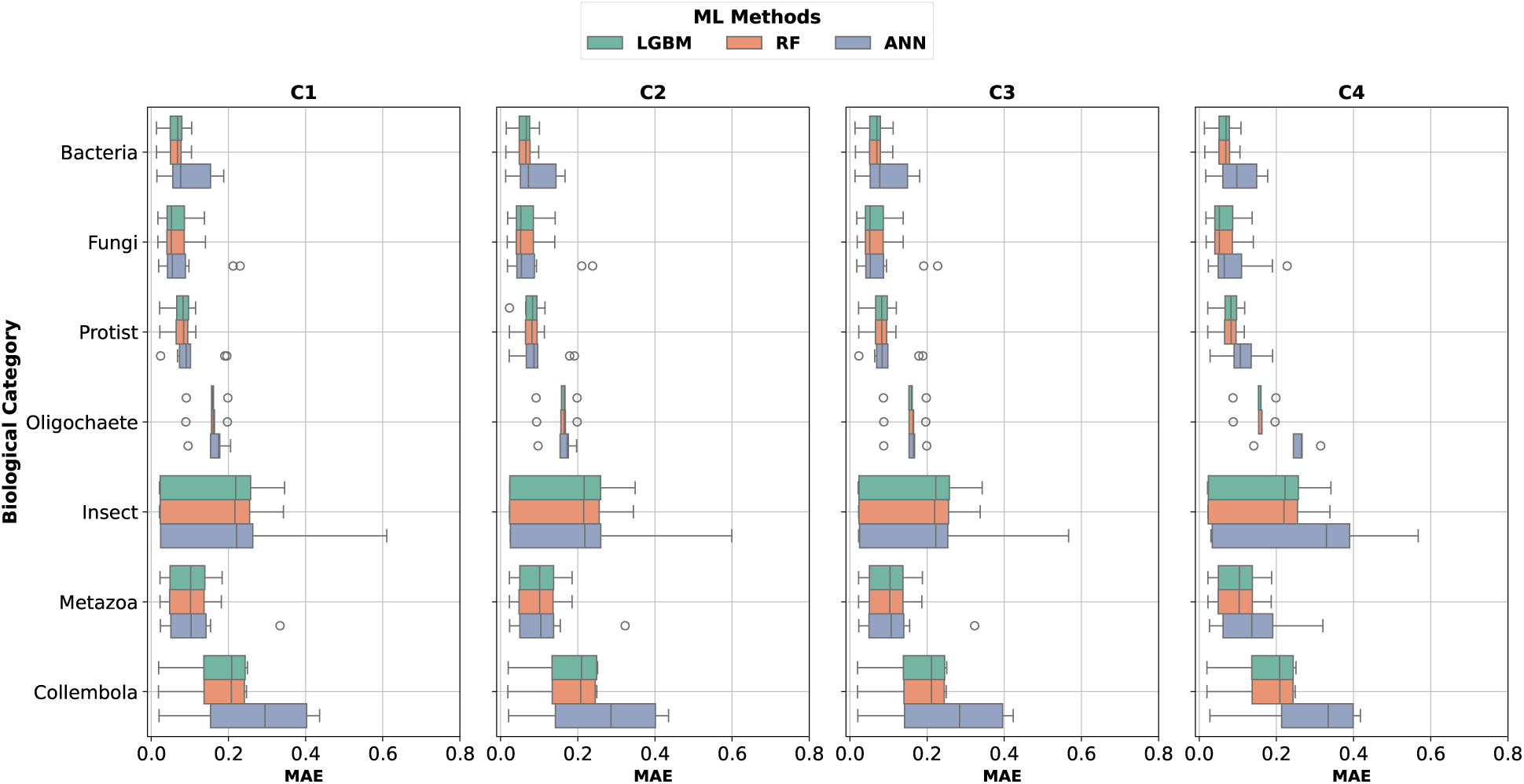
Comparison of model performance across trophic groups using MAE. The boxplots summarize the predictive performance for each taxon (Bacteria, Fungi, Protist, Oligochaete, Insect, Metazoa, and Collembola) across the four data configurations (C1–C4). Each configuration was evaluated using three machine learning models (LGBM, RF, and ANN).

**Figure B.11:**
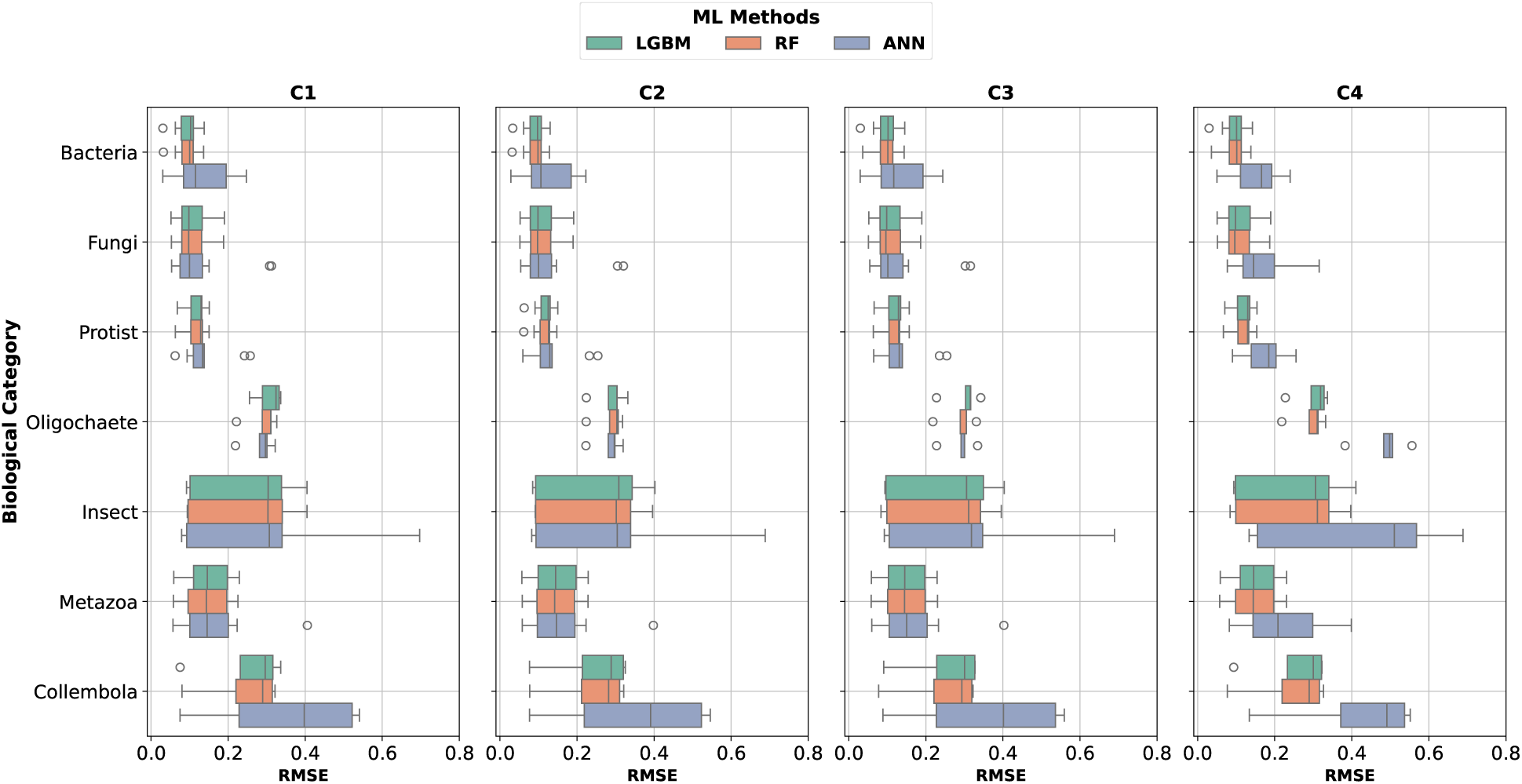
Comparison of model performance across trophic groups using RMSE. The boxplots summarize the predictive performance for each taxon (Bacteria, Fungi, Protist, Oligochaete, Insect, Metazoa, and Collembola) across the four data configurations (C1–C4). Each configuration was evaluated using three machine learning models (LGBM, RF, and ANN).

**Figure B.12:**
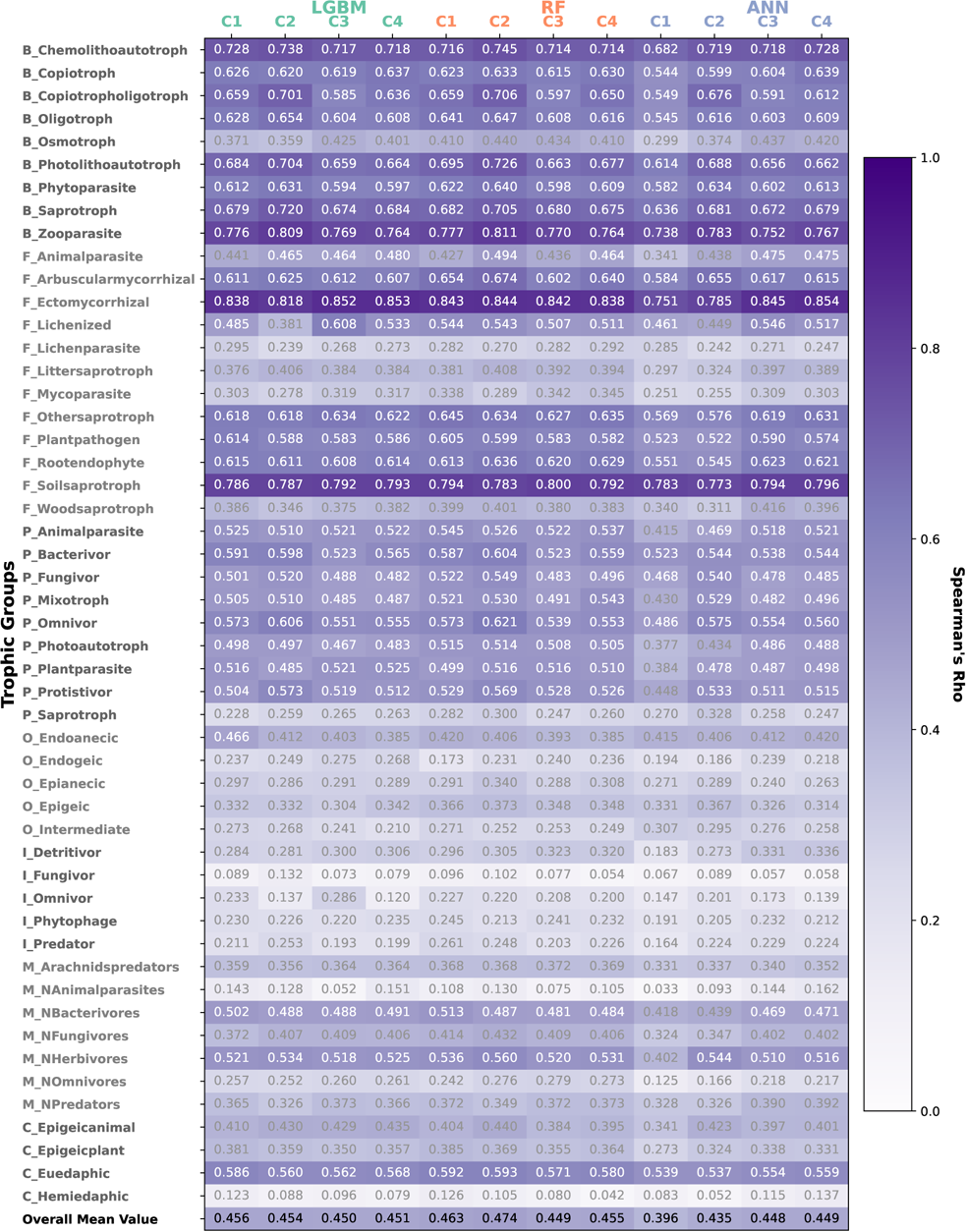
Heatmap of Spearman’s correlation for the best-performing models across the 51 trophic groups, ML techniques (LGBM, RF, ANN), and data configurations (C1–C4). The overall mean values at the bottom summarize model performance.

